# Dopamine promotes instrumental motivation, but reduces reward-related vigour

**DOI:** 10.1101/2020.03.30.010074

**Authors:** J.P. Grogan, T.R. Sandhu, M.T. Hu, S.G. Manohar

**Author notes:** Corresponding author =.

## Abstract

We can be motivated when reward depends on performance, or merely by the prospect of a guaranteed reward. Performance-dependent (contingent) reward is instrumental, relying on an internal action-outcome model, whereas motivation by guaranteed reward may serve to minimise opportunity cost in reward-rich environments. Competing theories propose that each type of motivation should be dependent on dopaminergic activity. We contrasted these two types of motivation with a rewarded saccade task, in patients with Parkinson’s disease (PD). When PD patients were ON dopamine, they had greater response vigour (peak saccadic velocity) for *contingent* rewards, whereas when PD patients were OFF medication, they had greater vigour for *guaranteed* rewards. These results support the view that reward expectation and contingency drive distinct motivational processes, and can be dissociated by manipulating dopaminergic activity. We posit that dopamine is necessary for goal-directed motivation, but dampens reward-driven vigour, challenging the theory that tonic dopamine encodes reward expectation.

## Introduction

Organisms expend more effort when their actions can lead to rewards, as the value of the reward offsets the extra effort expended to receive them (Kool & Botvinick, 2018; Manohar et al., 2015; Niv, Joel, & Dayan, 2006; Shenhav et al., 2017). They will even do so if the extra effort does not increase the reward they receive (Glaser et al., 2016; Milstein & Dorris, 2007; Xu-Wilson, Zee, & Shadmehr, 2009), indicating that mere expectation of reward is enough to justify the effort cost. Motivation, which promotes this effort expenditure, has two facets: it allows actions to be directed towards goals, and it energises our actions when rewards are expected (Niv et al., 2006). These two aspects are not always coupled. For example, employees might be salaried, where a fixed reward is *guaranteed* irrespective of achievements, or they might receive merit-based pay that is *contingent* on meeting performance targets (Lazear, 2000).

Motivation by contingent reward is instrumental, requiring an internal model of which outcome follows which action (Daw & Dayan, 2014; Dickinson, 1985). In contrast, motivation by increased reward expectation, independent of what an agent does, is model-free. One hypothesis is that this non-selective effect of incentives is adaptive, and is mediated by tonic dopamine (Niv, Daw, Joel, & Dayan, 2007). The rationale is that environmental rewards can vary, and it pays to capitalise on them when availability is high (Niv et al., 2007). For example, in a reward-rich environment, actions should be faster, because the opportunity cost of wasting time is higher (Shadmehr, De Xivry, Xu-Wilson, & Shih, 2010). Accordingly, when people merely expect more reward, they act faster – even if that speed does not increase their likelihood of reward – and may actually impair performance due to speed-accuracy trade-offs (Beierholm et al., 2013; Niv et al., 2007; Otto & Daw, 2019). Under this view, higher tonic dopamine levels should increase the opportunity cost of guaranteed rewards and thus amplify motivation (Beierholm et al., 2013; Niv et al., 2007), whereas dopamine would have little effect on motivation by contingent reward, which presumably relies on more sophisticated causal reasoning (Niv et al., 2006).

An alternative, contradictory prediction could be made from studies of model-based learning. Motivation by performance-contingent reward directs behaviour towards a goal, and might require an internal model of action-outcome-reward associations. Model-based learning requires dopamine transients (Sharpe et al., 2017), and is impaired when PD patients are withdrawn from their dopaminergic medication (Sharp, Foerde, Daw, & Shohamy, 2016), while model-free learning may be unaffected by dopamine (Wunderlich, Smittenaar, & Dolan, 2012). These studies suggest that dopamine is important for assigning reward within a causal model of the world, rather than directly to stimuli in a Pavlovian manner. If this were the case, then low-dopamine states should impair contingent motivation without affecting reward expectation effects. Guaranteed reward, in contrast, could motivate performance through simple reward associations, without causal understanding, and thus might not require dopamine.

Here we aim to test these two conflicting predictions, by asking whether dopamine affects motivation by performance-contingent reward, or motivation by expectation of guaranteed reward. We measured motor vigour in a monetary incentive saccade task in PD patients ON and OFF dopamine.

## Results

### Dopamine promotes contingent motivation and attenuates reward-expectation motivation

We tested 26 PD patients ON and OFF dopaminergic medication (PD ON & PD OFF) and 29 HC on a rewarded eye-movement task that separated effects of contingent and non-contingent motivation (see Fig.1a for task, see Table 2 for participant details). Participants made saccades to a target after hearing cues indicating how reward would be determined (Fig.1b). To measure motivation by contingent rewards, we compared trials where rewards were delivered depending on participants’ response times (Performance), to trials where rewards were given with 50% probability (Random). We matched the average reward rate so that both these conditions had identical reward expectation and uncertainty, and only differed in their *contingency*. To measure motivation by reward expectation, we compared trials with a guaranteed reward (10p) to those with a guaranteed no-reward (0p). In both these conditions rewards were delivered unconditionally, and only differed in terms of *expected reward*. In all trials, feedback was shown about whether the response was fast or slow, in addition to reward, to control for intrinsic motivation. As in previous work (Manohar, Finzi, Drew, & Husain, 2017), peak saccade velocity residuals (Fig.1c-e) provided the critical measure of response vigour. A three-way repeated-measures ANOVA tested whether dopamine differentially affected contingent and guaranteed motivation – manifest by a three-way (contingency*motivation*drug) interaction.

**Figure 1.**
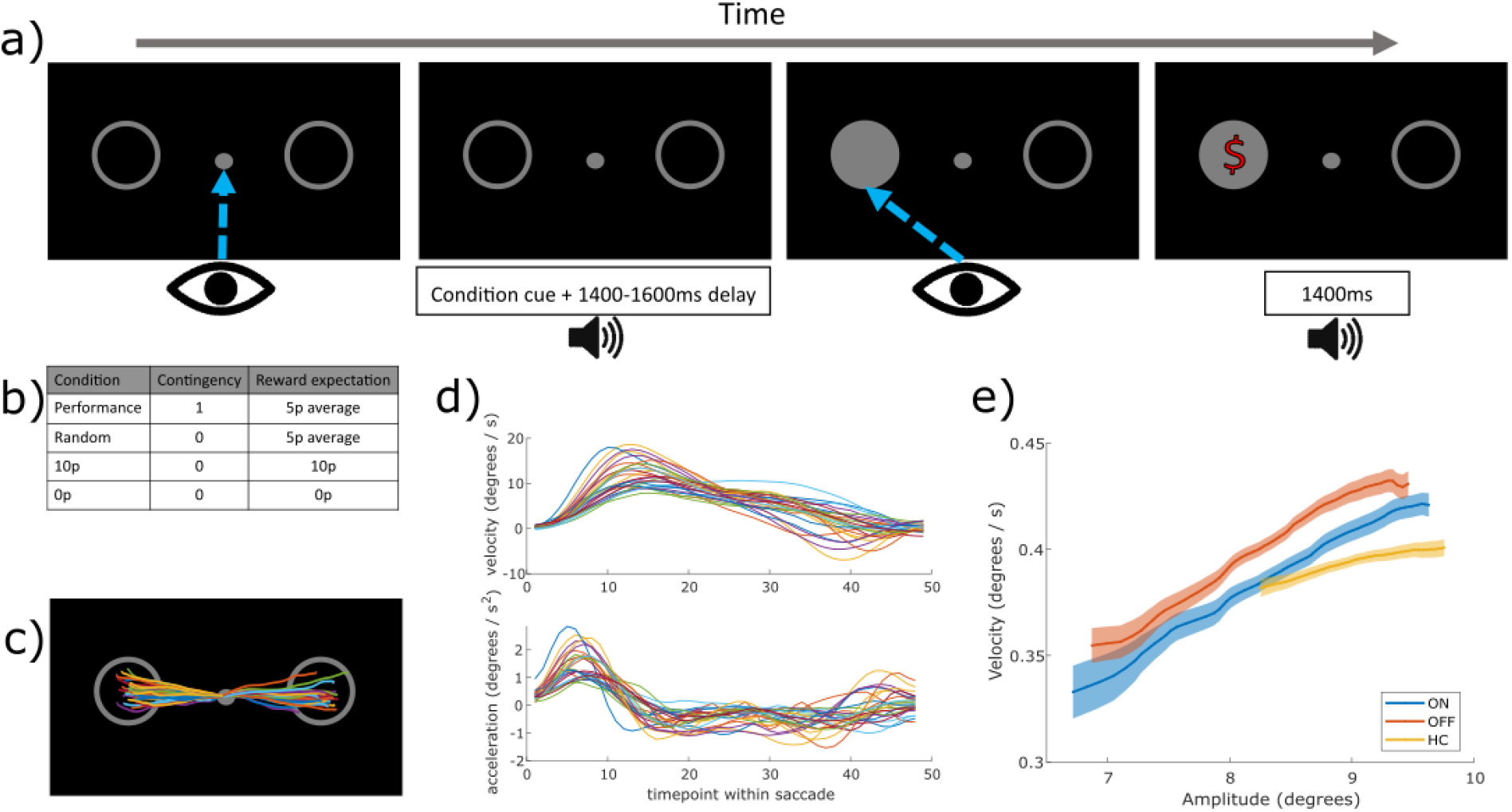
Saccade task design and example eye-tracking traces. a) Trial design: participants fixated on the centre, heard a cue for the condition (Performance/Random/10p/0p), waited a delay (1400/1500/1600ms) and then looked towards to the circle that lit up, and were given 10p or 0p reward depending on the condition, along with feedback on their response time (fast/slow). b) To measure contingent motivation, we compared “Performance” trials, where participants had to be faster than their median RT to win reward (thus giving 50% trials rewarded on average), with “Random” trials where a random 50% of trials were rewarded. To measure motivation by expected reward we compared “10p” trials where rewards were guaranteed, with “0p” trials where no-reward was guaranteed. c) Example eye-position traces for one participant and condition (different colours are different trials). d) Example mean velocity and acceleration profiles for all PD ON in the 10p condition. e) Peak velocity of individual saccades increase with the amplitude of movement – the “main sequence”; example showing the 10p condition, for PD ON, OFF and HC. Saccadic vigour, our measure of interest, was indexed by the residuals after regressing out amplitude from peak velocity, for each participant.

**Table 2.**
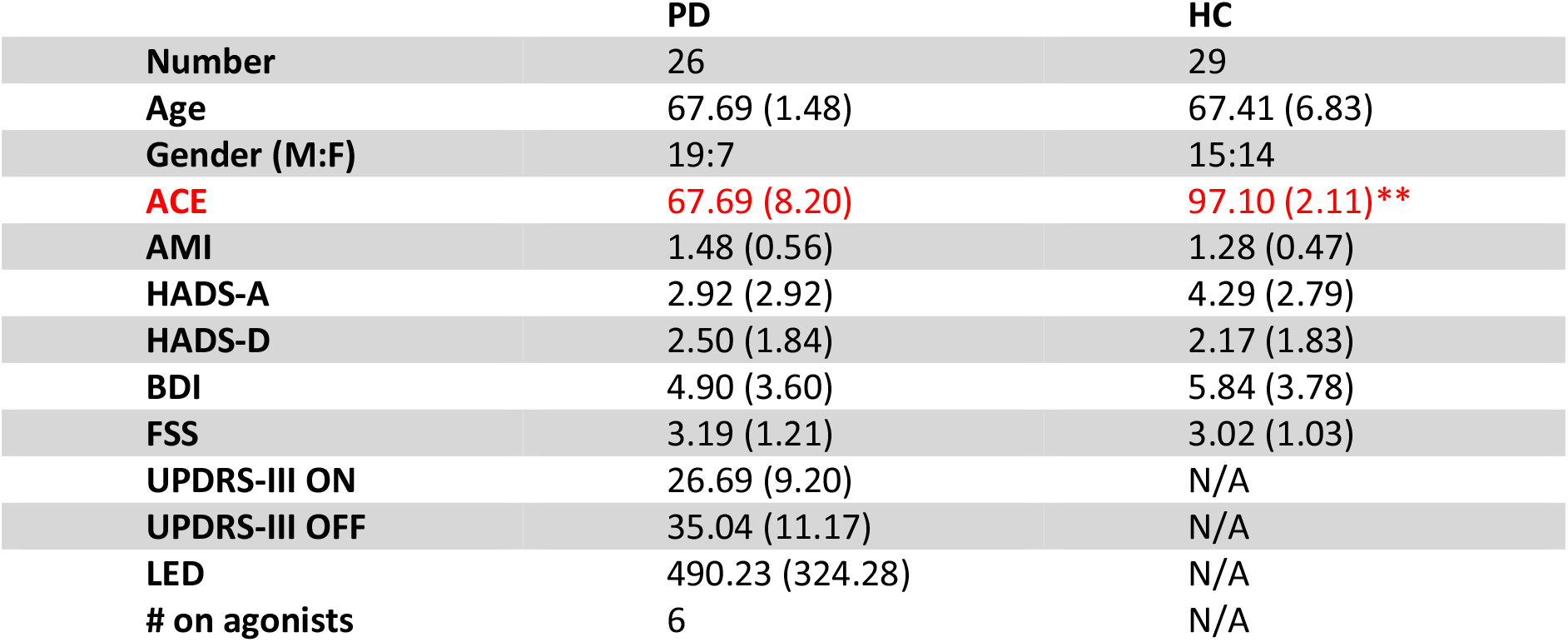
Participant demographics for PD patients and Healthy Controls (HC) included in the analysis. Standard deviations are given in parentheses. ** = p < .01 (independent samples t-test). ACE = Addenbrooke’s Cognitive Exam, AMI = Apathy & Motivation Index, HADS = Hospital Anxiety & Depression scores (A & D given separately), BDI = Beck Depression Inventory, FSS = Fatigue Severity Scale, UPDRS-III = Unified Parkinson’s disease rating scale Part 3, performed ON and OFF medication, LED = Daily Levodopa Equivalent Dose, # on agonists = number of patients taking dopamine agonists in addition to levodopa.

|  | PD | HC |
| --- | --- | --- |
| **Number** | 26 | 29 |
| **Age** | 67.69 (1.48) | 67.41 (6.83) |
| **Gender (M:F)** | 19:7 | 15:14 |
| **ACE** | 67.69 (8.20) | 97.10 (2.11)** |
| **AMI** | 1.48 (0.56) | 1.28 (0.47) |
| **HADS-A** | 2.92 (2.92) | 4.29 (2.79) |
| **HADS-D** | 2.50 (1.84) | 2.17 (1.83) |
| **BDI** | 4.90 (3.60) | 5.84 (3.78) |
| **FSS** | 3.19 (1.21) | 3.02 (1.03) |
| **UPDRS-III ON** | 26.69 (9.20) | N/A |
| **UPDRS-III OFF** | 35.04 (11.17) | N/A |
| **LED** | 490.23 (324.28) | N/A |
| **# on agonists** | 6 | N/A |

Dopaminergic medication significantly modulated how contingent and guaranteed motivation affected motor vigour (Fig.2a, three-way interaction on peak velocity residuals, p = .0023; see Table 1 for statistics). This was because, when ON medication, patients were motivated by contingency but not reward expectation (separate two-way ANOVA in PD ON: contingency*motivation, p = .0170; see Table S1, Fig.2b), whereas after overnight withdrawal of medication there was a borderline significant interaction in the opposite direction, as PD OFF were motivated by reward expectation but not contingency (PD OFF ANOVA: p = .0501; Table S1). This indicates that when PD patients were ON medication, motivation was strongest when reward was contingent on performance, but when they were OFF medication, patients were motivated by guaranteed rewards.

**Figure 2.**
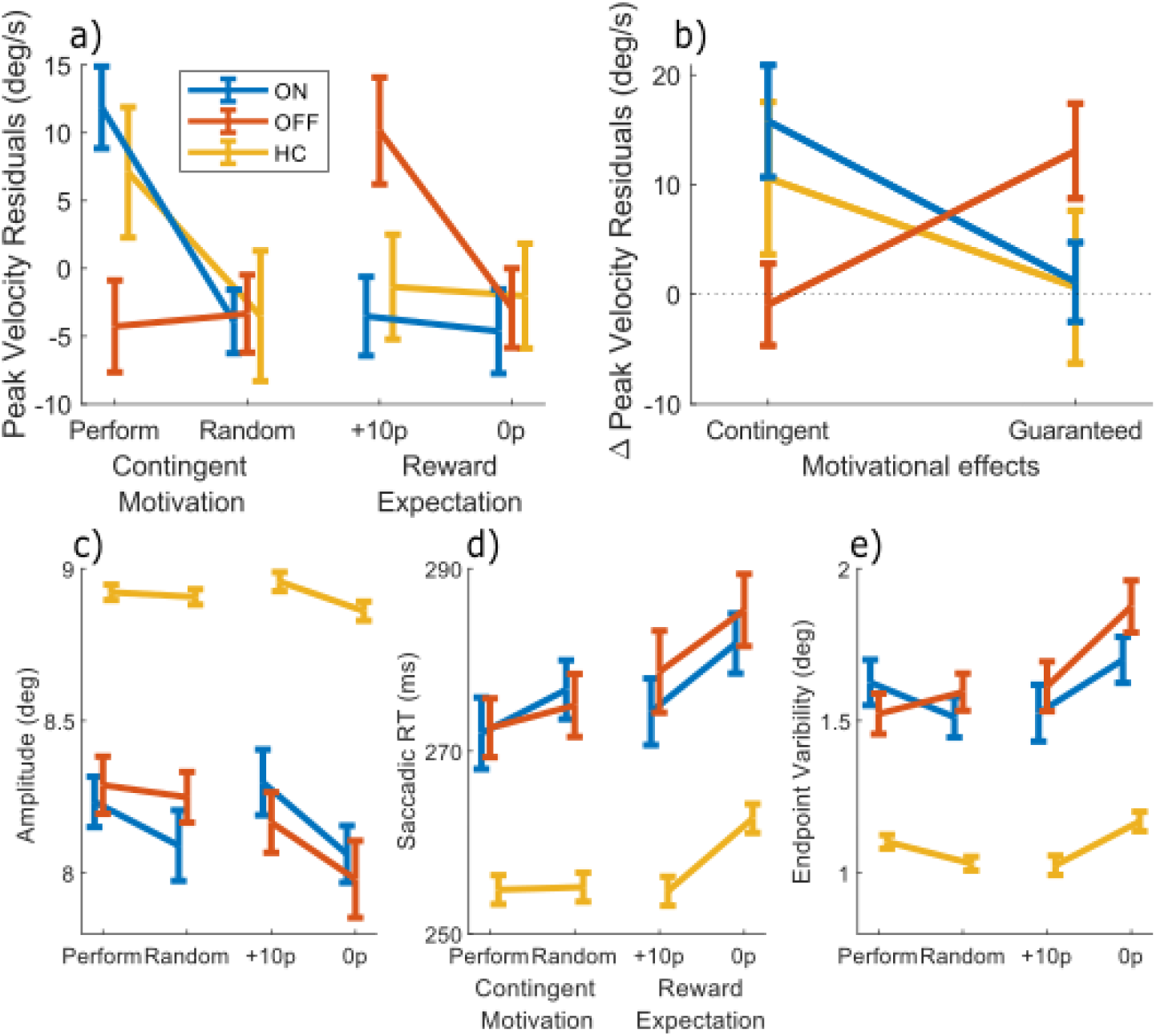
Differential effects of dopamine on two types of motivation. The mean measures for the four conditions (Performance, Random, Guaranteed 10p, Guaranteed 0p) for each variable. The difference between Performance and Random shows the effect of Contingent motivation, while the difference between 10p and 0p shows the motivating effect of Reward Expectation. a) Velocity residuals indexed behavioural vigour. When ON dopamine, patients were motivated to invigorate their saccades when reward depended on response time, but not when expecting a guaranteed reward. In contrast, when OFF dopamine, vigour was driven by expectation of guaranteed reward, but not by contingency. HC were similar to PD ON dopamine. b) The effects of contingent (Performance - Random) and guaranteed (10p - 0p) rewards on peak velocity. PD ON and HC have contingent invigoration, while PD OFF have guaranteed invigoration. c-e) No motivational effects were observed for c) saccade amplitude, d) saccade RT, or e) endpoint variability, although PD patients had slower, smaller and more variable saccades. All measures are in visual degrees, except saccade RT (ms). Error bars show within-subject SEM.

**Table 1.** Statistics for main behavioural analyses. Three-way (motivation*contingency*drug) repeated measures ANOVA on each behavioural measure, for the PD patients ON and OFF medication. An effect of contingency means the guaranteed conditions (10p, 0p) were different to the contingent conditions (Performance, Random). An effect of motivation means the 10p and Performance conditions were different to the Random and 0p conditions. An interaction of the two means that contingent rewards differed from guaranteed rewards. The Contingency * Motivation * Drug condition means that the effects of contingent and non-contingent rewards differed by PD medication state. Significant effects are highlighted in red. * = p < .05, ** = p < .01

| Measure | Effect | F (1, 200) | p | $\eta_p^2$ |
| --- | --- | --- | --- | --- |
| Velocity residuals | Motivation | 9.7704 | *.0020 | .0466 |
|  | Contingency | 0.0194 | .8895 | .0001 |
|  | Drug | 0.0004 | .9850 | .0000 |
|  | Motivation * Contingency | 0.0051 | .9429 | .0000 |
|  | Motivation * Drug | 0.2626 | .6089 | .0013 |
|  | Contingency * Drug | 11.1072 | ** .0010 | .0526 |
|  | Contingency * Motivation * Drug | 9.5190 | ** .0023 | .0454 |
| Amplitude | Motivation | 3.5577 | .0607 | .0175 |
|  | Contingency | 1.2284 | .2690 | .0061 |
|  | Drug | 0.0000 | .9984 | .0000 |
|  | Motivation * Contingency | 0.5545 | .4573 | .0028 |
|  | Motivation * Drug | 0.2278 | .6337 | .0011 |
|  | Contingency * Drug | 1.7763 | .1841 | .0088 |
|  | Contingency * Motivation * Drug | 0.0287 | .8655 | .0001 |
| Saccadic RT | Motivation | 3.4333 | .0654 | .0169 |
|  | Contingency | 4.2922 | *.0396 | .0210 |
|  | Drug | 0.3560 | .5514 | .0018 |
|  | Motivation * Contingency | 0.3663 | .5457 | .0018 |
|  | Motivation * Drug | 0.0694 | .7925 | .0003 |
|  | Contingency * Drug | 0.6246 | .4303 | .0031 |
|  | Contingency * Motivation * Drug | 0.0185 | .8920 | .0001 |
| Endpoint Variability | Motivation | 2.6780 | .1033 | .0132 |
|  | Contingency | 3.6181 | .0586 | .0178 |
|  | Drug | 1.0095 | .3162 | .0050 |
|  | Motivation * Contingency | 3.9524 | *.0482 | .0194 |
|  | Motivation * Drug | 1.2787 | .2595 | .0064 |
|  | Contingency * Drug | 1.3819 | .2412 | .0069 |
|  | Contingency * Motivation * Drug | 0.1626 | .6872 | .0008 |

To confirm that the effects on peak velocity residuals were not driven by changes in other aspects of saccades, the same 3-way ANOVA was run on each of the other saccade measures. Dopaminergic state did not affect amplitude or saccadic RT, although saccadic RT was quicker for both contingent conditions (p = .0396), and endpoint variability did have a contingency*motivation interaction (p = .0482, see Table 1, Fig2.c-e) as variability was higher for 0p condition.

We also compared both PD ON and OFF separately against the healthy age-matched controls (HC) with three-way mixed ANOVA, to see under which conditions patients deviated from healthy behaviour. HC had overall larger amplitudes, quicker saccadic RTs and lower endpoint variability than both PD ON or OFF (Fig.2a, see Table S3 and Table S4 for statistics). HC did not significantly differ from PD ON or OFF in velocity residuals, although their pattern was numerically closest to PD ON with greater contingent motivation.

When examining HC alone, they showed no significant effects or interactions on peak velocity (p > .05; see Table S1), although a motivation*contingency interaction was found on endpoint variability as contingent rewards increased variability while guaranteed rewards decreased it (p = .0048, see Table S2).

### Velocity profiles

The effects above demonstrate peak velocity shows strong effects of reward and dopamine, so next we examined the time-course of how velocity was modulated during a saccade. We computed the velocity across time within the movements, and compared the reward effects for PD ON and OFF using cluster-wise permutation tests. Contingent rewards (Performance – Random) did not significantly affect velocity or acceleration for PD ON or OFF, as permutation tests for each condition and the difference between conditions found no significant clusters (p > .05; Fig.3a). However, guaranteed rewards (10p – 0p) lead to greater velocity early in the saccade for PD OFF (p < .05; Fig.3d), which was significantly different from PD ON (p < .05). Acceleration traces showed this was due to PD OFF having greater acceleration early in the movement (Fig.3e, p < .05).

**Figure 3.**
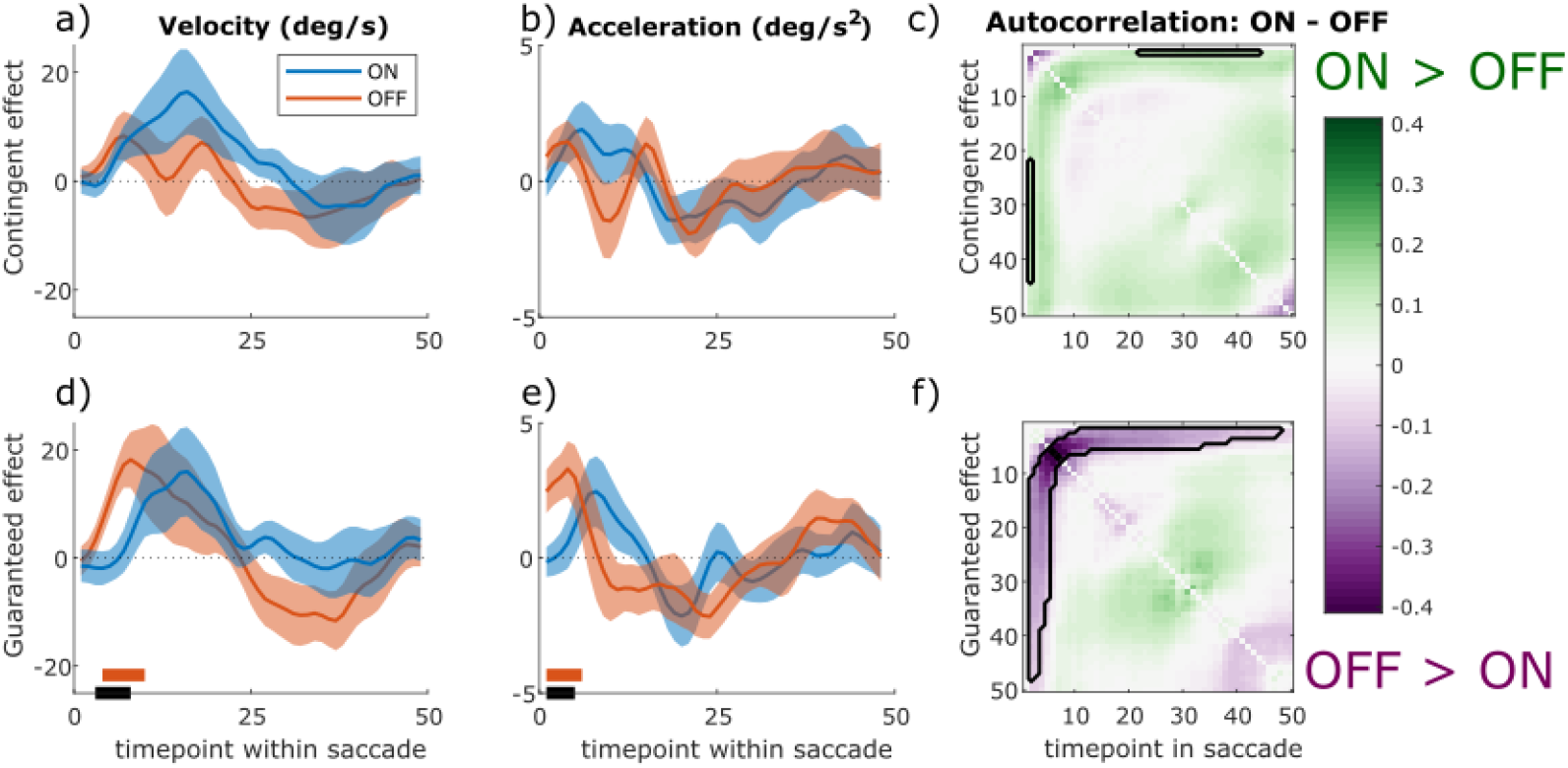
Motivational effects on instantaneous velocity, acceleration and eye-position autocorrelation within a saccade. The instantaneous velocity (a & d) is increased by contingent (a) and guaranteed (d) rewards, and PD patients OFF have an earlier and greater increase in velocity for guaranteed rewards than PD ON. The orange bar shows time-points where PD OFF had velocity significantly greater than zero (cluster-wise permutation tests, p < .05), the black bar shows time-points where PD ON and OFF significantly differed (PD ON did not differ from zero for any, so there is no blue bar). Shading shows SEM. Acceleration traces (b & e) showed this was due to guaranteed motivation increasing acceleration at the start of the movement for PD OFF (e; significant cluster, p < .05). The autocorrelation of eye position across the saccade (c & f) show that PD OFF have greater autocorrelation for guaranteed rewards early in the trial (f; purple colours = OFF > ON, black lines outline significant clusters), which is likely due to the greater velocity and acceleration increasing noise.

Faster movements are known to be more error-prone, but motivation can attenuate this effect, making movements more precise (Manohar, Muhammed, Fallon, & Husain, 2019). Autocorrelation of eye position over time within saccades provides an indicator of motivation improving precision. This improvement, provided by corrective motor signals, can be increased by incentives and is revealed by late reductions in autocorrelation (Codol, Holland, Manohar, & Galea, 2019). In the current study, guaranteed rewards led to greater autocorrelation early in the saccades for PD OFF than ON (shown as the decrease in Fig.3f). This is most likely because motivation increased acceleration, thus increasing noise in early parts of the movements. Notably, this reward-related correlation did not persist until the end of the saccades, suggesting that negative feedback corrected it. However, as we did not find decreased autocorrelation around the end of the saccades, this represents only indirect evidence of negative feedback.

### No correlation of the velocity effects for the distinct motivational processes

Previous work had shown that motivation by contingent and guaranteed reward did not correlate across participants (Manohar et al., 2017), so we asked whether dopamine’s effects upon these two types of motivation was also uncorrelated. We found no correlation between effects of contingent and guaranteed rewards on saccade velocity residuals in PD ON, PD OFF or HC separately, nor a correlation between medication states, nor between the drug-induced changes in the effects (p > .05; see Table S5 for statistics). This suggests that the two effects are separate and independent, and not antagonistic within the same person. In particular, the degree to which dopamine improved performance-contingent motivation did not predict the degree to which it reduced motivation by guaranteed rewards.

## Pupil dilatation

We examined pupil dilatation after the cue onset and before the target appeared (after 1400ms). Previous research has shown a greater effect of contingent than guaranteed reward on pupil dilatation, maximal around 1200ms after the cue (Manohar et al., 2017), so we used a window-of-interest analysis on the mean pupil dilatation 1000-1400ms after the cue. There were no significant effects or interactions (p > .05; Fig.5, see Table S6 for statistics), suggesting that dopamine and reward did not affect pupil responses in PD patients.

**Figure 5.**
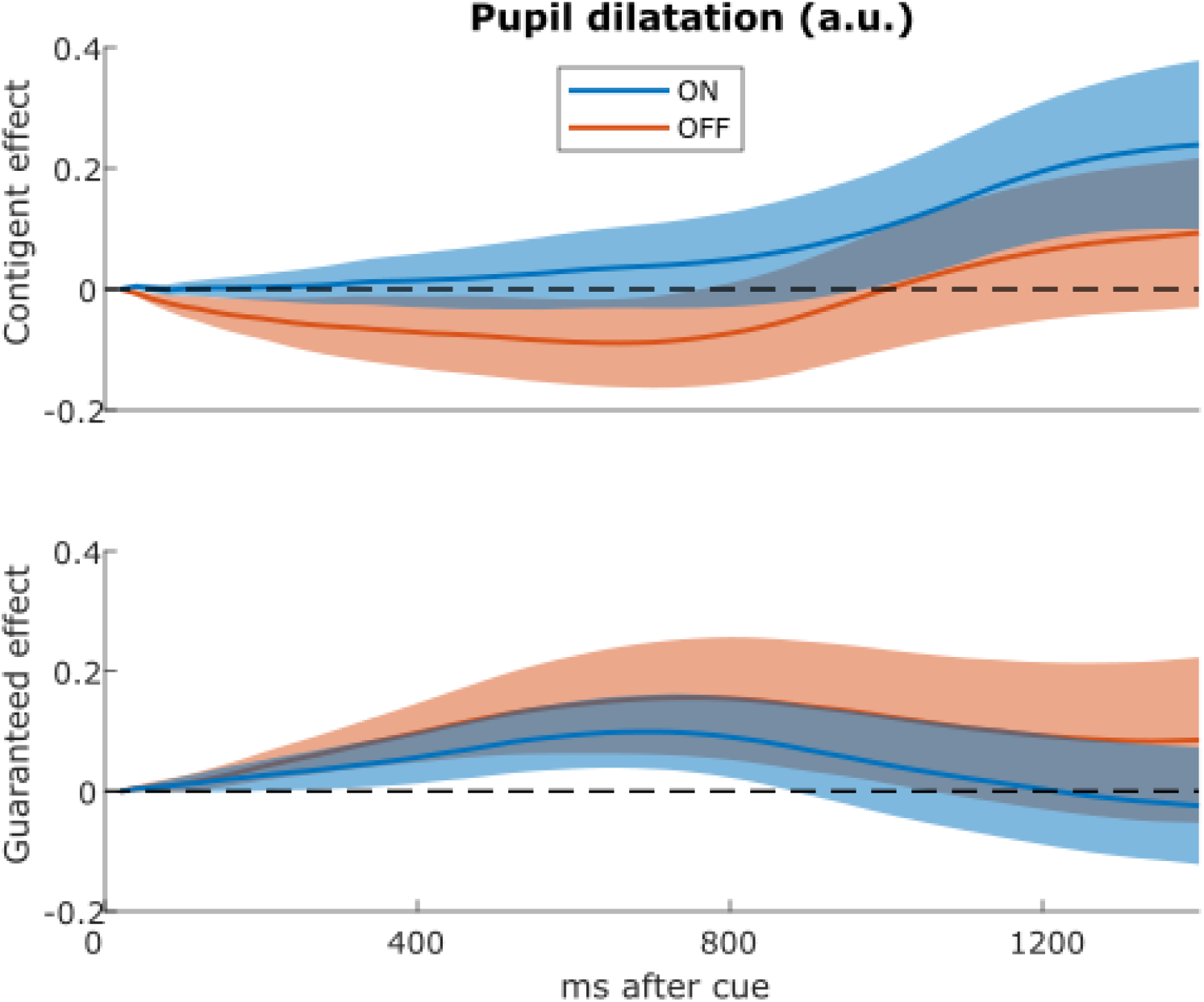
No effects of motivation on pupil dilatation. The effects of contingent (top) and guaranteed rewards (bottom) on pupil dilatation in the different conditions up to 1400ms after the reward cue. Pupil dilatation is baselined to the time of cue onset. There were no significant clusters of difference between PD ON and OFF (p > .05). Shading shows SEM.

We also used a hypothesis-free analysis, using cluster-wise permutation testing across the whole time course to look for significant differences between conditions, which also found no significant effects (p > .05).

We found no correlations between pupil dilatation and motivation effects in any group, or overall (p > .05; Table S7). Thus, the vigour effects were not related to pupillary dilatation before the movement.

## Discussion

In this study we tested two competing theories of dopaminergic motivation – that dopamine improves instrumental, contingent motivation, and that dopamine improves guaranteed reward motivation via reward expectation. Patients with PD made more vigorous responses, measured by peak saccade velocity residuals (Fig.2a), when rewards were either contingent on performance or guaranteed, but these two effects were differentially affected by dopaminergic medication. When ON medication, PD patients were motivated by rewards contingent on performance, but not by guaranteed rewards. In contrast, when patients were OFF their dopaminergic medication, the opposite pattern was observed; they were motivated by guaranteed rewards, but not by rewards contingent on performance. HC showed a similar pattern to PD ON. Guaranteed rewards led to PD OFF having earlier increases in velocity and acceleration (Fig.3b & d), which was not seen in PD ON or when rewards were contingent, and this was accompanied by increased autocorrelation of eye position (Fig.3f), suggesting increased motor noise early in the saccade. The motivational effects were uncorrelated across people and between medication states (Fig.4), indicating that dopamine does not promote one type of motivation over another in a competitive fashion, and were not associated with changes in pupil dilatation (Fig.5). Rather, reward expectation and contingency provide distinct motivational drives (Fig.6), which can be dissociated by dopaminergic medication.

**Figure 4.**
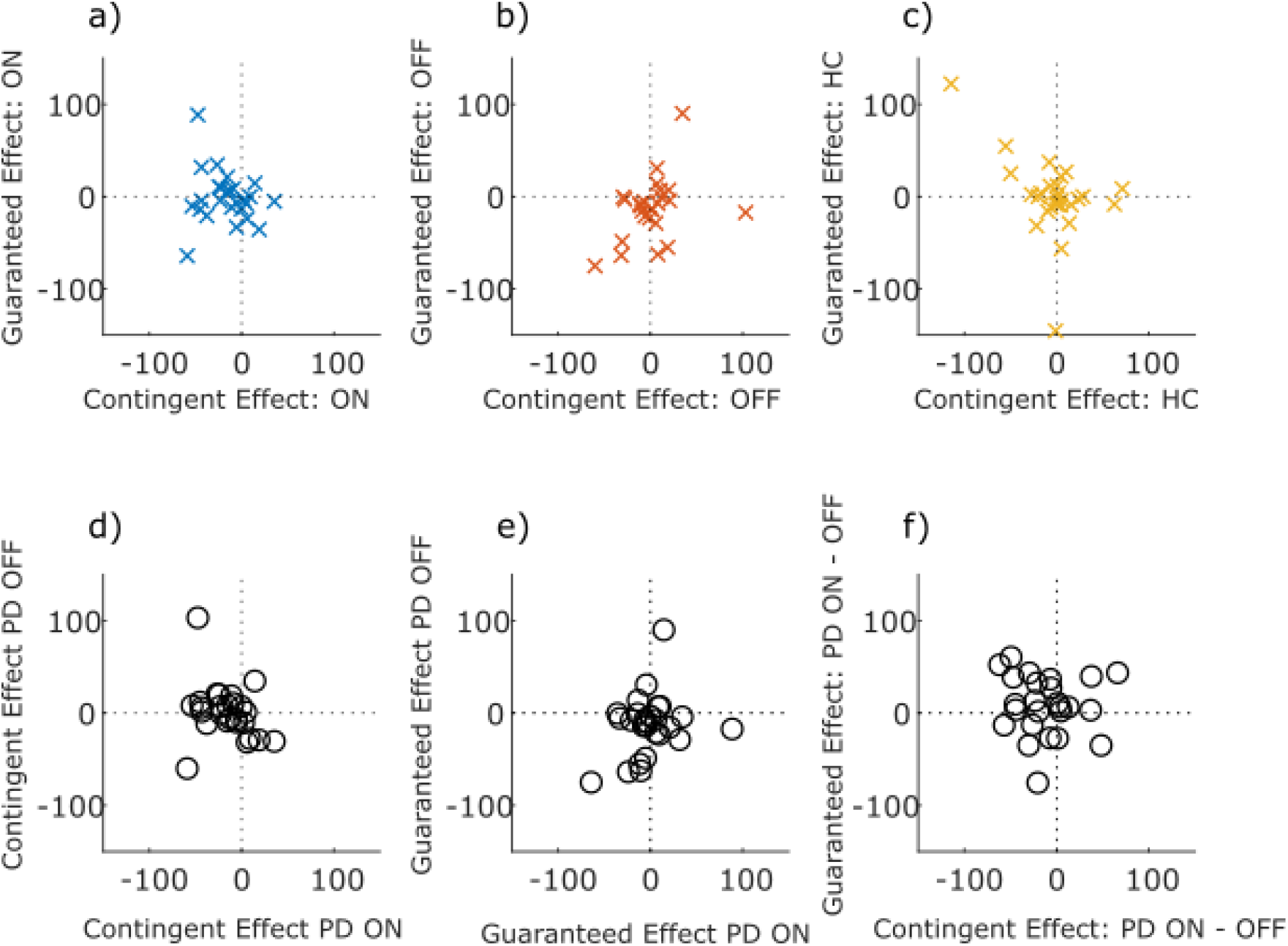
No correlations between contingent and guaranteed rewards. Scatter plots of the effect of contingent and guaranteed rewards (i.e. contingent effect = Performance minus Random trials, guaranteed effect = guaranteed 10p minus guaranteed 0p trials) on velocity residuals, within each group (top row: PD ON, PD OFF, HC), and between medication conditions (bottom row). No Spearman’s correlations were significant (p > .05; see Table S5 for statistics).

**Figure 6.**
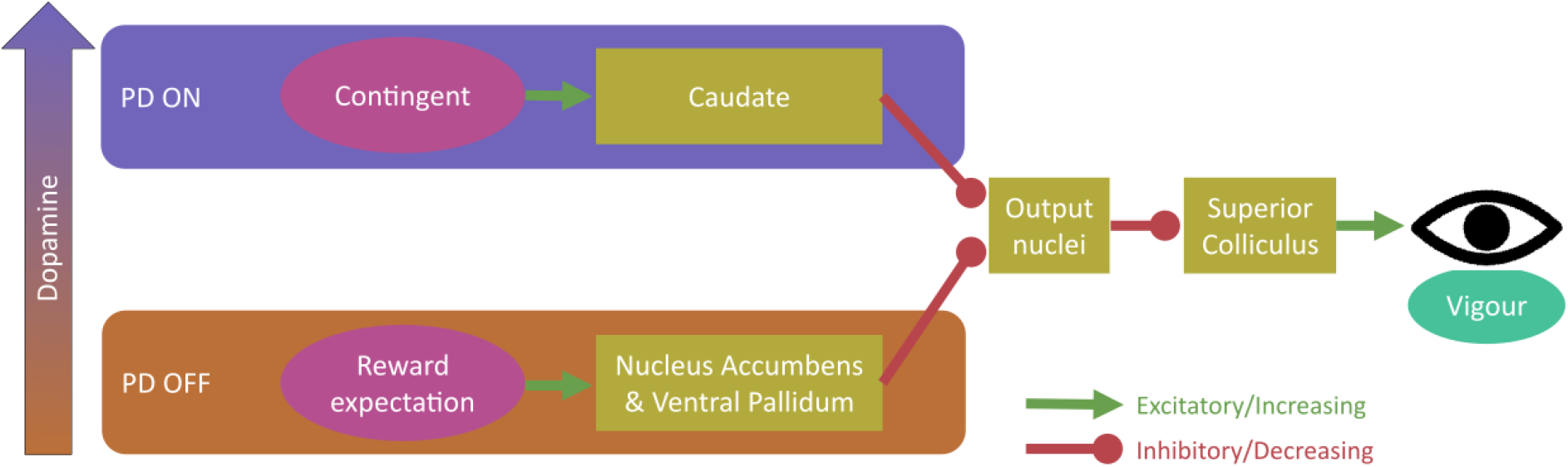
Proposed model for dopaminergic dissociation of reward expectation and contingent motivation. We propose that dopamine (in PD patients) increases contingent motivation by acting on the caudate nucleus, which disinhibits the superior colliculus (via the basal ganglia output nuclei) and affects the firing activity within the saccade, influencing vigour. Separately, high tonic dopamine impairs reward expectation motivation via the nucleus accumbens and ventral pallidum, which also disinhibit the basal ganglia output nuclei to affect superior colliculus firing activity and thus vigour within the saccade. Possible mechanism for this dissociative dopamine influence include separate dopaminergic regions innervating the two pathways, ‘global’ vs ‘local’ signalling, or different expression of D1-like and D2-like receptors (see text for details).

The results suggest that dopamine is necessary for contingent motivation. Contingent motivation requires an internal model of action-outcome-reward associations for goal-directed behaviour (Daw & Dayan, 2014; Dickinson, 1985), while reward expectation can occur via model-free stimulus-reward associations (Niv et al., 2007). While both model-free and model-based processes contribute to motor movements (Haith & Krakauer, 2013), these rely on separate neural circuits. Thus, the current work fits with previous research in which dopamine is necessary only for model-based, not model-free, learning (Sharp et al., 2016; Sharpe et al., 2017; Wunderlich et al., 2012).

Our finding that dopaminergic medication removes the reward expectation effect on vigour contrasts with previous research which suggests tonic dopamine levels encode the average reward rate, such that higher reward rates lead to greater dopaminergic tone, encouraging faster responding and greater vigour to minimise time spent without the expected reward – the opportunity cost (Beierholm et al., 2013; Niv et al., 2007). Our results show that reward expectation can still influence vigour when dopaminergic tone is low, yet does not when dopaminergic tone is high. It is possible that being ON dopamine led to tonic dopamine being so high that there was little difference between the high and low reward signals. However, this would have resulted in both high and low guaranteed reward conditions having high velocity when ON, comparable to PD OFF in the 10p condition, which was not seen. One explanation for the discrepancy with previous research is that the previous studies did not fully decouple contingent and non-contingent motivation, since the rewards were only given for successful performance (Beierholm et al., 2013; Niv et al., 2007), meaning the rewards were still contingent on performance. However, when separated, contingent motivation has larger effects on vigour than reward expectation (Manohar et al., 2017), and so it is possible that previously reported effects of average reward rate on vigour were due to the greater contingency, separate from or in addition to, reward expectation. An additional challenge to the tonic dopamine theory of reward expectation comes from the finding that fast phasic dopaminergic responses in the nucleus accumbens encode average reward rate, but slow tonic responses do not (Mohebi et al., 2019). That study suggests that reward expectation signals are independent of ventral tegmental area dopaminergic neuron firing, and may instead be due to ‘local’ control over nucleus accumbens core dopamine release. As dopamine is depleted in PD via dopamine-neuron death in the substantia nigra and ventral tegmental area, local dopamine release in other areas may be relatively preserved, and thus still able to influence vigour when PD patients are without dopamine.

The effect of reward-expectation on peak velocity was accompanied by greater velocity, acceleration, and autocorrelation early in the saccade for PD OFF than ON. Greater autocorrelation at this point is expected, as greater velocity increases noise (Harris & Wolpert, 2006; Paul M. Fitts, 1954). However, this noise increase did not persist until the end of the saccade, as there was no increase in autocorrelation at the end of the saccade (Fig3.f) and no greater endpoint variability (Fig2.d) – indeed, guaranteed rewards actually decreased endpoint variability, although this was not affected by dopamine. This offers some indirect evidence that the increased noise in this condition was attenuated via negative feedback (c.f. Manohar, Muhammed, Fallon, & Husain, 2019).

The age-matched HC showed a similar pattern to PD patients ON medication, but a different pattern to PD OFF, suggesting that dopaminergic medication may restore motivation to the ‘normal’ pattern for healthy older adults. However, the HC did not show any significant effects of motivation (contingent or reward-expectation) on peak velocity, saccadic RT, or amplitude, when analysed separately, despite showing effects on endpoint variability. This differs from the pattern previously found in healthy young adults, where reward expectation and contingent motivation both increase velocity (Manohar et al., 2017). Ageing might therefore decrease motivational influences on vigour, though direct comparison between these age groups in future work is needed to confirm this.

The motivational effects reported here were not related to any pupillary responses, unlike our previous findings in young people, which may be due to both ageing and PD decreasing the influence of rewards on pupil size (Manohar & Husain, 2015; Muhammed et al., 2016). Additionally, while the two distinct motivational effects on velocity were uncorrelated within PD patients, it is possible that subgroups of patients showed different effects. For example, whether patients were on D2 agonists (Bryce & Floresco, 2019) or had tremor-dominant disease (Wojtala et al., 2019) might be relevant. However this study was not powered to detect such differences as only 6 patients were taking agonists in addition to levodopa.

Considering the neuroanatomical differences between contingent motivation and reward expectation may help to explain our results. The nucleus accumbens and ventral pallidum modulate their activity by reward expectation (Mohebi et al., 2019; Tachibana & Hikosaka, 2012), while the caudate nucleus is active when rewards are contingent on behaviour (Tricomi, Delgado, & Fiez, 2004). Both the caudate and accumbens/pallidum project to the output nuclei of the basal ganglia, allowing saccade initiation via the superior colliculus, which controls not only the direction of saccades, but also their instantaneous velocity during the movement (Smalianchuk, Jagadisan, & Gandhi, 2018). We propose contingent motivation and reward expectation both lead to motivational signals affecting the superior colliculus’ activity controlling the velocity and acceleration of saccades, and these are differentially affected by dopamine (Fig.6), although we remain agnostic as to the mechanism for this difference. Possibilities include the two systems receiving input from separate regions of the dopaminergic system which are differentially depleted in PD (e.g. dopamine overdose hypothesis (Cools, 2006)), differences in “global” and “local” dopamine signals (Mohebi et al., 2019), or differences in D1-like and D2-like receptor expression within these systems (Surmeier, Ding, Day, Wang, & Shen, 2007; Yetnikoff, Lavezzi, Reichard, & Zahm, 2014). Further studies should address this question of the underlying mechanism.

We have shown that in PD, dopaminergic medication boosts motivation by contingent rewards, but reduces motivation by expected reward. Nonspecific invigoration by reward may thus be generated by a different neural system than goal-directed motivation. This suggests that dopaminergic medication may be a potential treatment for impairments in contingent motivation, but not for deficits related to reward expectation.

## Methods

### Participants

Thirty PD patients were recruited from volunteer databases in the University of Oxford. They were all taking levodopa medication, and some were also taking monoamine oxidase inhibitors and/or dopamine agonists (Table 2). They were randomly assigned to be tested ON or OFF medication first, and withdrawn from standard release medication for 16+ hours and controlled-release medication for 24+ hours. Two patients did not complete both sessions, and two did not have enough trials that passed all the criteria (see Analysis section) so were excluded, leaving 26 patients. Thirty healthy controls (HC) were recruited from volunteer databases also, and tested once, and one HC was excluded for insufficient trials passing the criteria.

All participants gave written informed consent, and ethical approval was granted by the South Central Oxford A REC (18/SC/0448).

### Procedure

The task was run in Matlab (www.mathworks.com, version 7) using the Psychophysics toolbox (Kleiner et al., 2007), on a Windows XP computer with a CRT monitor (1024×768 pixels, 40×30cm, 100Hz refresh rate) at 70cm viewing distance. Eye movements and pupil size were recorded with Eyelink1000 at 1000Hz.

On each trial of the task a fixation dot (0.3° radius) was presented at the centre of the screen, with two empty circles (1.1° radius) shown 9.3° to the left and right of the fixation dot. After 500ms of fixation, a cue was given by a voice over the speaker, indicating the type of trial the participant was in:

- “Performance” indicated that fast response times would win 10p, while slow response times would win 0p
- “Random” indicated a 50% probability of 10p or 0p, regardless of response time
- “Ten pence” indicated a guaranteed 10p, regardless of response time
- “Zero pence” indicated guaranteed 0p, regardless of response time

A delay of 1400, 1500 or 1600ms was given (with equal probability), after which one of the two circles turned white (50% probability of left or right) and participants had to saccade to this circle to complete the trial and receive the outcome.

Participants could only affect the outcome in the Performance condition (by moving faster); all others were independent of their speed. In the Performance condition, rewards were based upon response time (i.e. total time between the target appearing and gaze arriving at the target), which is only minimally influenced by saccade velocity. Participants were rewarded when response time was quicker than their recent median response time for the last 20 Performance trials, which thus yielded a 50% reward rate overall. The Random condition acts as a control to these trials, with a random 50% of trials rewarded, and thus equal expected value but with no performance-contingency. Rewards in the guaranteed conditions also had zero contingency on performance, but yielded different expected rewards (10p vs 0p), thus comparing them indexes the pure effect of expecting reward. Notably, participants always received feedback on their speed (fast/slow, using median split over 20 previous trials in that condition), regardless of reward, to minimise differences in intrinsic motivation.

There were 12 types of each trial in a block, in a random order, and participants completed 4 blocks. The blocks differed in the modality of feedback; blocks giving auditory feedback on the speed and visual feedback on the reward, and vice versa for the other two blocks. This order was randomised across participants.

### Analysis

The Performance and 10p conditions are high motivation conditions. The difference between Performance and Random conditions gives the effect of contingent motivation, while the difference between 10p and 0p conditions gives the effect of reward expectation.

As in previous studies (Manohar et al., 2017), our primary measure of interest was saccadic vigour. We measured peak saccade velocity on each trial. We took the first saccade after target onset which was greater than 1° in amplitude, and used a sliding window of 4ms width to calculate velocity, excluding segments faster than 3000°s^−1^ or where eye tracking was lost. Saccades with peak velocities outside 80-2500°s^−1^ were excluded, as were trials where participants reached the target before 180ms or after 580ms. Two PD patients and one HC had fewer than 10 trials that passed these criteria for one condition, so were excluded from the analysis.

To remove the main sequence effect of amplitude on velocity (Harris & Wolpert, 2006), we regressed velocity against amplitude and took the residual velocity as our measure of interest. This was done for each participant’s separate session. We also measured amplitude, saccadic reaction time (RT), and endpoint variability of these saccades. Saccadic RT is the time between the target onset and the start of the saccade.

To analyse velocity and acceleration traces, and autocorrelation and covariance of the eye movements we linearly interpolated 50 points along each saccade to move them into the same units. Instantaneous velocity was smoothed across 3 time-points, while acceleration was smoothed across 5. We also calculated velocity and acceleration traces on the raw (non-interpolated) traces and then interpolated them afterwards, which gave very similar results.

Pupil dilatation was measured in arbitrary units (a.u.) relative to the baseline pupil size at the cue onset. Blinks under 500ms were linearly interpolated, steps in pupil size above 2.5 a.u./ms were removed, and data were averaged in 20ms bins for plotting.

We used `rmanova` from the matlib toolbox (https://github.com/sgmanohar/matlib) to perform analyses – this uses `fitglme` to perform the repeated measures test and anova to perform hypothesis tests on the GLME. We used three-way repeated measures ANOVA to compare effects of motivation, contingency and dopaminergic medication in PD patients, and followed this up with two-way ANOVA when a three-way interaction was found. To compare each PD condition against HC we used mixed ANOVA. We also used cluster-wise permutation tests for the time course data (velocity, acceleration, pupil dilatation, autocorrelation and covariance), to control the family-wise error rate at .05.

## Data and Code Availability

Analyses were performed in Matlab using custom scripts, which are available on GitHub (https://doi.org/10.5281/zenodo.3786452). Anonymous data are available on OSF (https://osf.io/2k6x3), as is the experiment file (osf.io/y9xhp).

## Acknowledgements

We would like to thank the patients and participants for their time in taking part in this study, and the funders for their support (MRC Clinician Scientist Fellowship to SGM, MR/P00878X).

## Author Contributions

SGM designed the study, MH contributed to the patient cohort and manuscript feedback, TRS and JPG tested the participants, JPG analysed the data, and JPG wrote the manuscript with feedback from TRS and SGM.

