## Supplementary for "Dopamine promotes instrumental motivation, but reduces reward-related vigour"

### Supplementary Materials

#### Individual data on main measures

Here we plot the individual data for Fig.2.

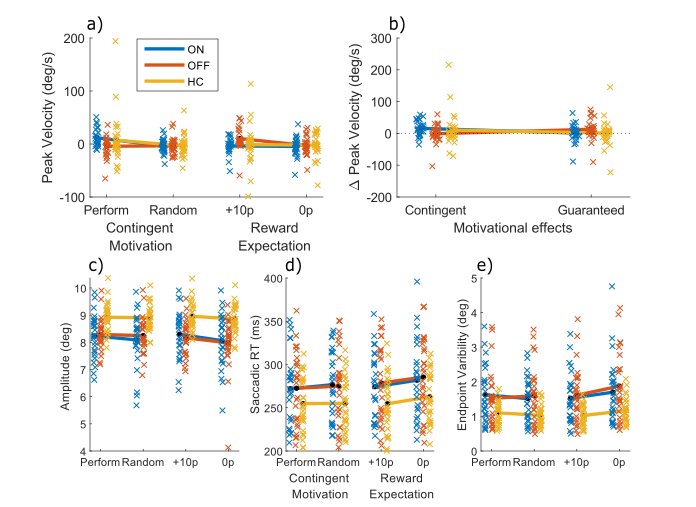

Figure S1. Individual data for saccadic measures. Crosses show each individual’s mean score for each condition, while black dots and lines show the means for each group for the four conditions (Performance, Random, Guaranteed 10p, Guaranteed 0p) for each variable: a) Velocity residuals, b) saccadic amplitude, c) saccadic RT, d) saccadic endpoint variability. All measures are in visual degrees, except saccade RT (ms).

#### Effects of motivation and contingency in separate groups

We ran separate two-way, repeated-measures ANOVAs on PD ON and PD OFF (and HC), to see how the three-way ANOVA (see main text, Table 1) broke down.

Table S1. Statistics for two-way ANOVA on peak velocity residuals for each group, separately. PD ON had effects of motivation, contingency and a trend interaction of these (as motivation increased peak velocity only for the contingent condition). PD OFF had no overall effect of contingency (as peak velocity was higher for guaranteed conditions) and a borderline significant motivation*contingency interaction (as 10p had the highest velocity). HC had no significant effects or interactions. * = p < .05, ** = p < .01.

| Group | Effect | F | p | $\boldsymbol{\eta}_{\boldsymbol{p}}^{\boldsymbol{2}}$ |
| --- | --- | --- | --- | --- |
| PD ON (*df* = 1, 100) | Motivation | 7.8283 | **.0062 | .0726 |
|  | Contingency | 7.1289 | **.0089 | .0665 |
|  | Motivation * Contingency | 5.8942 | *.0170 | .0557 |
| PD OFF (*df* = 1, 100) | Motivation | 2.9575 | .0886 | .0287 |
|  | Contingency | 4.4168 | *.0381 | .0423 |
|  | Motivation * Contingency | 3.9330 | .0501 | .0378 |
| HC (*df* = 1, 112) | Motivation | 0.9019 | .3443 | .0080 |
|  | Contingency | 0.3463 | .5574 | .0031 |
|  | Motivation * Contingency | 0.6995 | .4047 | .0062 |

We also ran two-way repeated-measures ANOVAs on saccade amplitude, RT, and endpoint variability in HC.

Table S2. Statistics for two-way ANOVA on other saccade measures in HC. HC had a motivation*contingency interaction for endpoint variability, as contingent rewards increased variability while expected rewards decreased it. ** = p < .01.

| Group | Effect | F (*df* = 1, 112) | p | $\boldsymbol{\eta}_{\boldsymbol{p}}^{\boldsymbol{2}}$ |
| --- | --- | --- | --- | --- |
| Amplitude | Motivation | 2.3510 | .1280 | .0206 |
|  | Contingency | 0.0255 | .8734 | .0002 |
|  | Motivation * Contingency | 1.2551 | .2650 | .0111 |
| Saccade RT | Motivation | 3.2227 | .0753 | .0280 |
|  | Contingency | 2.5743 | .1114 | .0225 |
|  | Motivation * Contingency | 2.7992 | .0971 | .0244 |
| Endpoint Variability | Motivation | 0.9304 | .3368 | .0082 |
|  | Contingency | 0.6651 | .4165 | .0059 |
|  | Motivation * Contingency | 8.2781 | **.0048 | .0688 |

#### Effects of PD on saccade measures

We ran separate mixed ANOVAs to compare HC against PD ON and PD OFF, and did this for each saccade measure separately. Here we present the statistics of those tests.

Table S3. Statistics for HC vs PD ON behavioural analyses. PD ON did not differ in velocity residuals from HC. There was a significant motivation*contingency interaction for endpoint variability, as both HC and PD ON showed greater variability after contingent rewards, but less variability after expected rewards. HC had greater amplitudes, quicker RTs and smaller variabilities than PD ON. * = p < .05, *** = p < .001

| Measure | Effect | F (*df* = 1, 220) | p | $\boldsymbol{\eta}_{\boldsymbol{p}}^{\boldsymbol{2}}$ |
| --- | --- | --- | --- | --- |
| Velocity residuals | Motivation | 4.1837 | *.0421 | .0194 |
|  | Contingency | 2.8162 | .0948 | .0131 |
|  | Group | 0.0008 | .9769 | .0000 |
|  | Motivation * Contingency | 3.1877 | .0756 | .0148 |
|  | Motivation * Group | 0.1677 | .6826 | .0008 |
|  | Contingency * Group | 0.4415 | .5071 | .0021 |
|  | Contingency * Motivation * Group | 0.1188 | .7306 | .0006 |
| Amplitude | Motivation | 2.5358 | .1128 | .0118 |
|  | Contingency | 0.0062 | .9371 | .0000 |
|  | Group | 85.0115 | ***<.001 | .2862 |
|  | Motivation * Contingency | 0.3183 | .5732 | .0015 |
|  | Motivation * Group | 0.7524 | .3867 | .0035 |
|  | Contingency * Group | 0.0239 | .8773 | .0001 |
|  | Contingency * Motivation * Group | 0.0012 | .9726 | .0000 |
| Saccade RT | Motivation | 1.6851 | .1957 | .0079 |
|  | Contingency | 0.8771 | .3501 | .0041 |
|  | Group | 23.0563 | ***<.001 | .0981 |
|  | Motivation * Contingency | 0.4339 | .5108 | .0020 |
|  | Motivation * Group | 0.0645 | .7998 | .0003 |
|  | Contingency * Group | 0.0000 | .9974 | .0000 |
|  | Contingency * Motivation * Group | 0.0989 | .7535 | .0005 |
| End-point Variability | Motivation | 0.3487 | .5555 | .0016 |
|  | Contingency | 0.4345 | .5105 | .0020 |
|  | Group | 55.8485 | ***<.001 | .2085 |
|  | Motivation * Contingency | 4.9883 | * .0266 | .0230 |
|  | Motivation * Group | 0.0025 | .9601 | .0000 |
|  | Contingency * Group | 0.0138 | .9066 | .0001 |
|  | Contingency * Motivation * Group | 0.1040 | .7474 | .0005 |

Table S4. Statistics for HC vs PD OFF behavioural analyses. HC did not significantly differ from PD OFF in velocity residuals, but HC had greater amplitudes, quicker RTs and lower variability than PD OFF. * = p < .05, *** = p < .001

| Measure | Effect | F (*df* = 1, 220) | p | $\boldsymbol{\eta}_{\boldsymbol{p}}^{\boldsymbol{2}}$ |
| --- | --- | --- | --- | --- |
| Velocity residuals | Motivation | 2.7173 | .1007 | .0127 |
|  | Contingency | 0.3062 | .5806 | .0014 |
|  | Group | 0.0016 | .9678 | .0000 |
|  | Motivation * Contingency | 0.0826 | .7741 | .0004 |
|  | Motivation * Group | 0.0038 | .9508 | .0000 |
|  | Contingency * Group | 2.3610 | .1259 | .0110 |
|  | Contingency * Motivation * Group | 2.8383 | .0935 | .0132 |
| Amplitude | Motivation | 1.3011 | .2553 | .0061 |
|  | Contingency | 1.8546 | .1747 | .0087 |
|  | Group | 89.4129 | ***<.001 | .2966 |
|  | Motivation * Contingency | 0.5945 | 0.4415 | .0028 |
|  | Motivation * Group | 0.1489 | 0.6999 | .0007 |
|  | Contingency * Group | 1.6466 | 0.2008 | .0077 |
|  | Contingency * Motivation * Group | 0.0482 | 0.8264 | .0002 |
| Saccade RT | Motivation | 1.3548 | 0.2458 | .0063 |
|  | Contingency | 2.5505 | 0.1117 | .0119 |
|  | Group | 30.5031 | ***<.001 | .1258 |
|  | Motivation * Contingency | 0.6381 | 0.4253 | .0030 |
|  | Motivation * Group | 0.0041 | 0.9490 | .0000 |
|  | Contingency * Group | 0.3769 | 0.5399 | .0018 |
|  | Contingency * Motivation * Group | 0.0512 | 0.8212 | .0002 |
| End-point Variability | Motivation | 2.6523 | 0.1049 | .0124 |
|  | Contingency | 3.0066 | 0.0844 | .0140 |
|  | Group | 62.8641 | ***<.001 | .2287 |
|  | Motivation * Contingency | 2.6619 | 0.1043 | .0124 |
|  | Motivation * Group | 1.0998 | 0.2955 | .0052 |
|  | Contingency * Group | 1.5467 | 0.2150 | .0072 |
|  | Contingency * Motivation * Group | 0.0096 | 0.9219 | .0000 |

#### Correlation of motivational effects

Here we present the statistical outputs of the Spearman’s correlations between motivational effects on velocity residuals within people and across drug states.

Table S5. Correlation statistics for motivational effects on velocity residuals. The Spearman’s rho and p-values for each pair of correlations (Fig.4) are shown here. The correlations were run on the effects: contingent effect = Performance minus Random; Guaranteed effect = guaranteed 10p minus guaranteed 0p; drug effects = ON minus OFF. No correlations were significant.

| Measures | Rho | p |
| --- | --- | --- |
| PD ON: Contingent effect vs Guaranteed effect | -0.1549 | .4503 |
| PD OFF: Contingent effect vs Guaranteed effect | 0.3730 | .0614 |
| HC: Contingent effect vs Guaranteed effect | -0.2153 | .2609 |
| Contingent effect: PD ON vs OFF | -0.3429 | .0869 |
| Guaranteed effect: PD ON vs OFF | 0.1432 | .4834 |
| Drug effects: Contingent effect vs Certain effect | -0.2438 | .2291 |

#### Pupil effects

*Table S6.* Pupil dilatation Window-of-interest statistics. A repeated-measures ANOVA comparing PD ON vs OFF on the mean pupil dilatation (from baseline) in the window-of-interest (1000-1400ms after the cue). There were no significant effects or interactions.

| Effect | F (*df* = 1, 216) | p | $\boldsymbol{\eta}_{\boldsymbol{p}}^{\boldsymbol{2}}$ |
| --- | --- | --- | --- |
| Motivation | 0.3525 | 0.5533 | .0016 |
| Contingency | 0.2743 | 0.6010 | .0013 |
| Drug | 1.4265 | 0.2336 | .0066 |
| Motivation * Contingency | 0.3106 | 0.5779 | .0014 |
| Motivation * Drug | 0.0331 | 0.8558 | .0002 |
| Contingency * Drug | 0.2589 | 0.6114 | .0012 |
| Motivation * Contingency * Drug | 0.3029 | 0.5826 | .0014 |

Table S7. No correlations between motivational effects on pupil dilatation and velocity residuals. Spearman’s correlation coefficients and p-values for correlations of the effects of contingent and guaranteed motivation on mean pupil dilatation change from 1000-1400ms after cue onset, and the effects on velocity residuals. No correlations were significant.

| Group | Motivation effect | Rho | P |
| --- | --- | --- | --- |
| ON | Contingent | -0.2759 | .1720 |
|  | Guaranteed | -0.3224 | .1085 |
| OFF | Contingent | -0.0182 | .9257 |
|  | Guaranteed | -0.0813 | .6931 |
| HC | Contingent | -0.2123 | .2964 |
|  | Guaranteed | -0.0990 | .6080 |

#### Eye position autocorrelations and covariance

Examining the time-time covariances and correlations of eye position during saccades can reveal effects of feedback or noise during movements. For example, reduced covariance towards the end of movements indicates corrective negative feedback. This has previously shown that people increase their feedback during rewarded movements to give more accurate saccades (Manohar, Muhammed, Fallon, & Husain, 2019). As we found that PD OFF had greater peak velocity, and an earlier increase in velocity (Fig.2a, Fig.3d), and greater velocities correspond to greater noise, examining the covariance can show whether feedback was also modulated to ameliorate the effects on accuracy.

Guaranteed motivation increased the (signed log) covariance for PD OFF early in the saccade (cluster-permutation testing, p < .05, outlined area in Fig.S2). This is around the time when PD patients showed the increased velocity in the saccade. As this increase did not persist until later in the saccade, it implies feedback corrected the extra noise caused by the greater velocity.

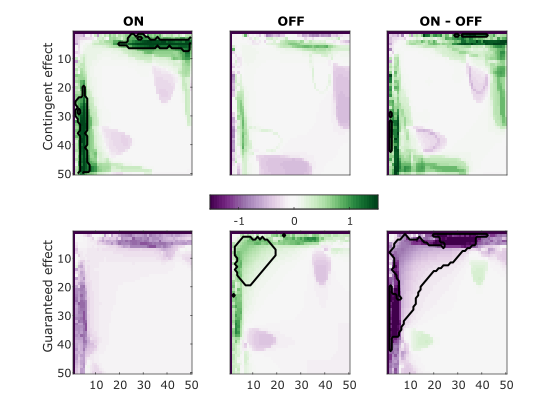

Figure S2. Motivational effects on time-time covariance within saccades. Each plot shows the effect of motivation (top row = contingent, bottom row = guaranteed) on the signed log covariance matrix, for PD ON (left), OFF (middle) and the difference between them (right). Green means the effect increased covariance, purple means a decrease, and the significant clusters are outlined in black. Cluster-wise permutation testing found that PD OFF had greater variance early on when rewards were guaranteed.

Comparing the drug effects (ON – OFF) on contingent and guaranteed motivation revealed that contingency significantly increased variance during a very small portion of the saccade more in PD ON than OFF, while guaranteed motivation increased variance more in PD OFF than ON early in the saccade (corresponding to a reduction for ON-OFF; Fig.S2).

We also examined the auto-correlations of the movements; correlation controls for the overall variance of the movements across trials, thus any differences seen in the covariance but not correlation may be due to differences in variance between conditions or groups.

We compared the Fisher transformed correlation matrices between conditions and groups with cluster permutation testing. PD OFF had a period of higher correlation early in saccades when rewards were guaranteed, and similar to the covariance. This effect was significant when compared against PD ON. PD ON also had a small period of increased autocorrelation midway through the saccade similar to the covariance, albeit a small cluster.

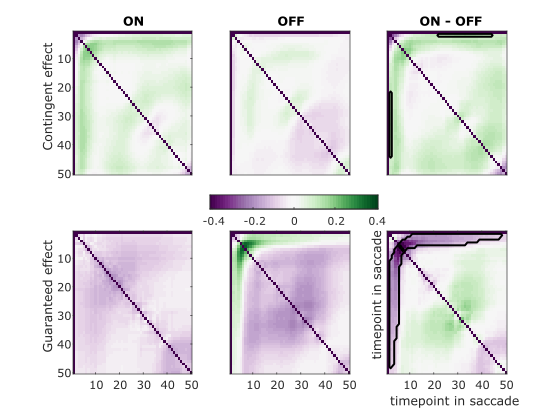

Figure S3. Motivational effects on saccade autocorrelation. The difference in (Fisher transformed) autocorrelation matrices for the contingent vs random (top row), and guaranteed reward vs no-reward (bottom row), for PD ON (left) and OFF (middle) and the difference (ON-OFF; right). Green means the motivational effect increased autocorrelation, purple means a decrease, black lines show significant clusters of difference. PD OFF have greater autocorrelation early in the saccades when rewards are guaranteed.

Taken together, the covariance and correlations show that reward expectation increased eye position variance early in the saccade for PD OFF. PD patients OFF had an earlier increase in velocity within a saccade (Fig.3d), which would increase variance also (Harris & Wolpert, 2006). However, this increased variance did not persist to the end of the movement, indicating feedback corrected it. This could also explain the lack of increased endpoint variability in these saccades despite the higher velocity. While we did not see differences in the correlations and covariance at the end of the saccade, this is indirect evidence of increased negative feedback on eye trajectories when motivated.

#### UPDRS

We looked to see whether the dopaminergic effects could be tied to PD symptom expression. The UPDRS is a measure of PD symptom severity and was performed in each session; part III measures motor symptom severity. We found no correlations between UPDRS scores and reward effects on velocities in PD ON or OFF (p > .05). Thus the reward effects were unrelated to PD symptom severity.

Table S8. No correlations between UPDRS-III scores and velocity residuals. Spearman’s correlation coefficients and p-values for correlation between effects of contingent and certain rewards on velocity residuals, and UPDRS scores, ON and OFF meds. No correlations were significant.

| PD Group | Effect | rho | p |
| --- | --- | --- | --- |
| ON | Contingent | -0.1256 | .5410 |
|  | Guaranteed | -0.2327 | .2527 |
| OFF | Contingent | -0.2067 | .3110 |
|  | Guaranteed | 0.1553 | .4487 |

#### Fixation period

We looked at whether motivation was affecting behaviour during the fixation period differently, which could potentially lead to differences during the movements. We excluded trials with saccades, blinks, deviations greater than 1.8° and segments with velocities greater than 30°s^-1^. We Fourier transformed the eye positions during the early (200-700ms after cue) and late (700-1200ms) fixation periods and compared these between conditions. We also looked at microsaccade (<1°) frequency, and ocular drift speed during the fixation period.

Here, we present the comparisons of PD ON and OFF; PD OFF had more microsaccades than PD ON (F (1, 201) = 5.0451, p = .0258) but no other effects or interactions were significant (p > .05).

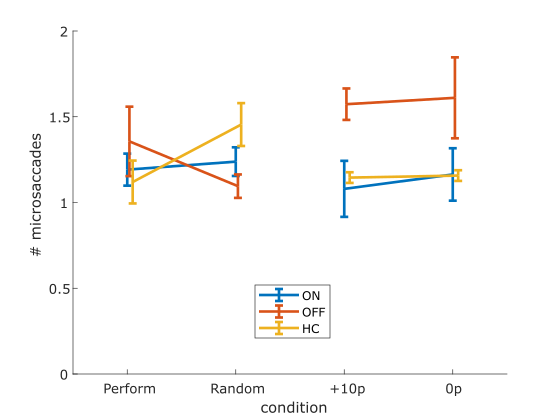

Figure S4. Number of microsaccades during the fixation period. PD OFF had more microsaccades overall, but there were no effects of condition or interactions. Error bars show SEM.

Similarly, ocular drift velocity differed between drug states (F (1, 216) = 5.4327, p = .0207), but not by condition (p > .05).

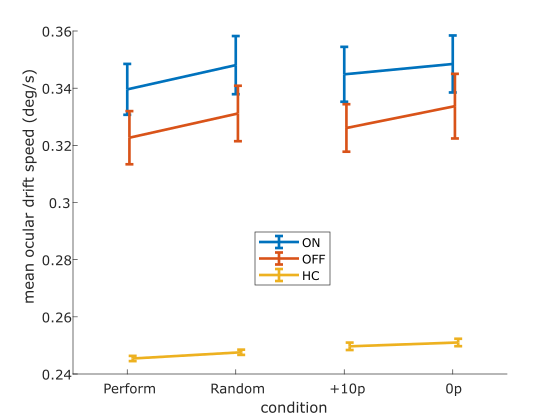

Figure S5. Mean eye drift speed during fixation period. HC had lower drift speeds than PD patients, and PD ON had higher drift speeds than OFF, but there were no effects of condition or interactions. Error bars are SEM.

To quantify ocular tremor, we performed Fourier transforms on the eye position in the early (200-700ms) and late (700-1200ms) fixation periods, which did not differ between drug states (p > .05).

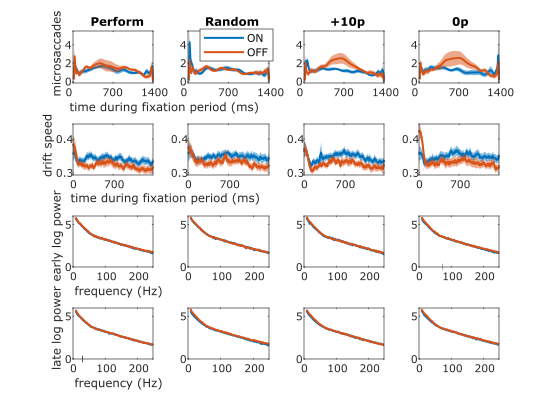

Figure S6. Fixation period measures for each condition. The microsaccade density (top) and mean speed (second row) across the fixation period, and the Fourier power spectra for the early (third row) and late (bottom) section of the fixation period, for each condition. There were no significant clusters of difference. Shading shows SEM.

Below, the reward effects for contingent and guaranteed rewards are shown for the same measures (10ms smoothing window used for microsaccades and drift speed, 20Hz smoothing window for power spectra). There were no significant differences (permutation clustering) between PD ON and OFF.

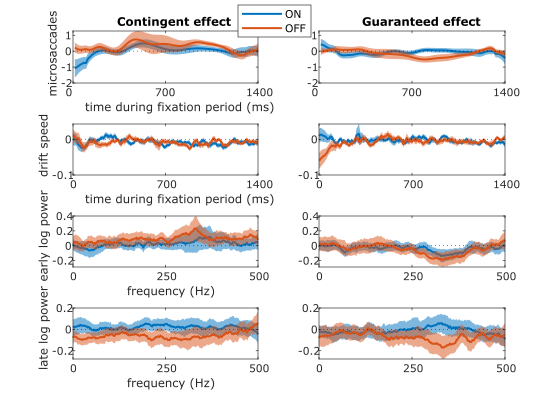

Figure S7. Motivation effects on the fixation period measures. The difference between contingent and random conditions (left), and guaranteed reward and non-reward (right), for the microsaccade density, mean drift speed, early and late period power spectra. Microsaccade density and mean ocular drift speed were smoothed with a 10ms moving window, the power spectra with 20Hz moving window. There were no significant clusters of difference. Shading shows SEM.
